# Regenerative capacity of neural tissue scales with changes in tissue mechanics post injury

**DOI:** 10.1101/2022.12.12.517822

**Authors:** Alejandro Carnicer-Lombarte, Damiano G. Barone, James W. Fawcett, Kristian Franze

**Affiliations:** John van Geest Centre for Brain Repair, Department of Clinical Neurosciences, University of Cambridge, Cambridge CB2 0PY, UK; Department of Physiology, Development and Neuroscience, University of Cambridge, Cambridge CB2 3DY, UK; Electrical Engineering Division, Department of Engineering, University of Cambridge, Cambridge CB3 0FA, UK; Centre for Reconstructive Neuroscience, Institute for Experimental Medicine CAS, Prague, Czech Republic; Institute of Medical Physics and Micro-Tissue Engineering, Friedrich-Alexander Universität Erlangen-Nürnberg, 91052 Erlangen, Germany; Max-Planck-Zentrum für Physik und Medizin, 91054 Erlangen, Germany

**Keywords:** Nerve stiffness, nerve injury, Crush injury, nerve compartments, tissue mechanics, Schwann cells

## Abstract

Spinal cord injuries have devastating consequences for humans, as mammalian neurons of the central nervous system (CNS) cannot regenerate. In the peripheral nervous system (PNS), however, neurons may regenerate to restore lost function following injury. While mammalian CNS tissue softens after injury, how PNS tissue mechanics changes in response to mechanical trauma is currently poorly understood. Here we characterised mechanical rat nerve tissue properties before and after *in vivo* crush and transection injuries using atomic force microscopy-based indentation measurements. Unlike CNS tissue, PNS tissue significantly stiffened after both types of tissue damage, likely mainly due to an increase in collagen I levels. Schwann cells, which crucially support PNS regeneration, became more motile and proliferative on stiffer substrates *in vitro*, suggesting that changes in tissue stiffness may play a key role in facilitating or impeding nervous system regeneration.

## Introduction

Traumatic injuries to the nervous system lead to axon damage and loss of neuronal signal conduction, resulting in loss of function as areas of the body become disconnected from the brain. When an injury occurs in the mammalian central nervous system (CNS), scar tissue develops which, while necessary to limit further damage to nearby tissue, also inhibits regrowth of axons through the lesion site. In contrast, injury to the peripheral nervous system (PNS) is followed by a robust regenerative response during which axons may regrow and reconnect with their targets^1^.

This opposite response of the CNS and PNS to injury has been subject of intense research, focusing largely on the chemical makeup of the injured tissue. Key molecules identified include factors inhibiting axon regeneration in the CNS^2–4^ and factors enhancing regeneration after PNS injuries^1^. However, the manipulation of molecular components in the CNS has so far failed to fully reproduce the robust regrowth of axons seen following PNS injuries^4^, indicating that an important component is still missing.

Injuries are not only accompanied by changes in a tissue’s chemical landscape but also in the mechanical microenvironment to which cells are exposed^5–7^. Tissue stiffness has recently been shown to regulate many cellular processes *in vivo*, including axon growth^8,9^and cell migration^10^. Also inflammation in the nervous system is triggered by non-physiological mechanical signals^11–14^, suggesting that changes in tissue mechanics may impact neuronal regeneration. In contrast to most other tissue types, which stiffen during the scarring process following an injury^15,16^, scars in the mammalian CNS are softer than healthy brain or spinal cord tissue^5,6^. However, whether the observed softening of neural tissue post injury is a general feature of the nervous system or if it is specific to the non-regenerative mammalian CNS^5^ is currently unclear.

We here characterised the stiffness of the various compartments of peripheral rat nerve tissue at the cellular scale using atomic force microscopy (AFM). With an apparent reduced elastic modulus *K* < 100 Pa, all nerve compartments were very soft and similar in stiffness to healthy brain and spinal cord white matter^5,17,18^. However, in contrast to CNS tissue^5,6^, nerve tissue stiffened in response to both crush and transection injuries, predominantly correlating with an increase in collagen I levels. Furthermore, proliferation and migration rates of primary Schwann cells – the main cell type promoting axon regeneration following peripheral nerve injury^19,20^ – increased significantly on substrates of increasing stiffness, suggesting that changes in neural tissue stiffness post injury may indeed contribute to regulating neuronal regeneration in both the mammalian CNS and PNS – in opposite directions.

## Results

### Mechanical properties of nerve endoneurium, perineurium, and epineurium

Nerves are composed of a variety of cell types and extracellular matrix (ECM) proteins, organised in a hierarchical structure. The nerve endoneurium, which makes up most of the nerve cross-sectional area, houses all neuronal axons and Schwann cells. Sections of endoneurium are bundled into fascicles by tightly packed perineurial cells, which make up the nerve perineurium. Finally, the outer portion of the nerve is protected by the ECM-rich epineurium consisting of fibroblast-like cells and collagen, which is subdivided into an outer epifascicular and an inner interfascicular portion^21^ (Fig 1A). We investigated the mechanical stiffness of these four nerve compartments in acutely excised *ex vivo* sciatic nerve preparations using AFM indentation measurements, in which forces were applied at similar length (∼μm), time (∼s), and force (∼nN) scales at which cells probe their environment^22^.

**Fig 1.**
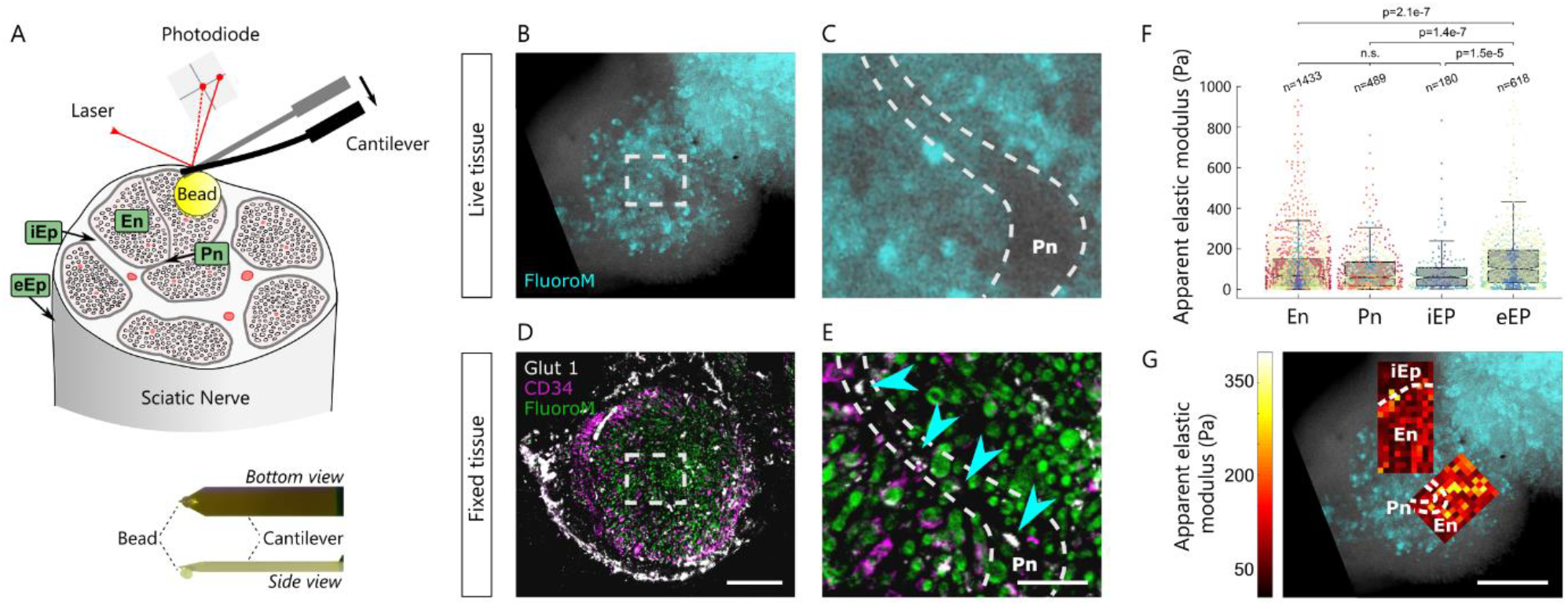
*Ex vivo* mechanical characterisation of nerve compartments. (A) Diagram of testing setup. A rat sciatic nerve cross-section exposes the various nerve compartments: endoneurium (En), perineurium (Pn), interfascicular epineurium (iEp), and epifascicular epineurium (eEp), which are measured by AFM.. (B-E) Images of nerve cross-sections. (B-C) Live imaging of fresh tissue, stained with Fluoromyelin live stain (cyan) to identify the different nerve compartments (indicated by white dashed lines in (C) and (E)). (B) Image as seen during AFM measurements. (C) Magnification of the area indicated by the white rectangle in (B). (D-E) The same tissue fixed and immunostained for myelin by Fluoromyelin (green), fibroblasts by CD34 (magenta), and perineurial cells by Glut1 (white) after the AFM measurements. (E) Magnification of the perineurium. The perineurium is identified as a thin strip of tissue lacking Fluoromyelin stain in live tissue (C, area between dashed lines), or containing Glut1+ perineurial cells in fixed and stained tissue (E, indicated by cyan arrows and dashed lines). The wider areas of the perineurium are visible in live imaging, and AFM measurements of the perineurium are performed over this region. Scale bars: 200 μm (B,D) and 50 μm (C,EF). (F) Box and scatter plots of tissue stiffness values for nerve compartments. All nerve compartments shared a similarly low stiffness with a median *K* ∼ 60 Pa, except for the significantly stiffer epifascicular epineurium (median *K* ∼ 101 Pa, p < 0.001, Kruskal-Wallis and Dunn-Sidak multiple comparisons test). Measurements *n* represented by individual dots, with measurements taken from the same animal depicted with the same colour. En: *N*=16 animals, *n*=1433 measurements, Pn: *N*=17, *n*=489, iEp: *N*=4, *n*=180, eEp: *N*=6, *n*=618. n.s.: not significant (all multiple comparison p values > 0.05). (G) Region in (B) with overlaid stiffness map recorded via AFM, colour codes local apparent elastic modulus *K*. Scale bar: 200 μm.

To identify the nerve compartments for AFM analysis, we stained cross-sections of nerve samples from the sciatic nerve of adult rats with the live marker fluoromyelin, which labels myelin sheets around axons in the endoneurium and highlights both the unlabelled narrow perineurium and the thicker interfascicular epineurium (Fig 1B-C). Immunostaining of fixed nerve tissue for markers of perineurial cells (Glut 1) and nerve fibroblasts commonly found in the epineurium (CD34)^23^ confirmed that fluoromyelin provided an accurate guide to identify nerve compartments in live nerve samples (Fig 1D-E). The interfascicular epineurium was identified as the tissue enveloping the exterior of the endoneurial cross-sections, and the epifascicular epineurium accessed by placing an un-sectioned sample of nerve on its side.

We performed AFM-based raster scans in areas of nerve cross-sections containing all compartments and pooled the measurements belonging to each nerve compartment for each nerve (Fig 1F-G). The elastic stiffness of endoneurium, perineurium, and interfascicular epineurium was similar with median apparent reduced elastic moduli *K* = 66 Pa, 59 Pa, and 59 Pa, respectively. The epifascicular epineurium was significantly stiffer than the other nerve tissues (median *K* = 101 Pa, *p* < 0.001, Kruskal-Wallis and Dunn-Sidak multiple comparisons test). All samples had a similarly wide spread of data, with some measurements yielding values of up to *K* ∼ 900 Pa, indicating a large degree of heterogeneity.

While technically not feasible for perineurium and epineurium, nerve endoneurium was significantly stiffer when probed over longitudinal sections if compared to cross-sections (median *K* = 120 Pa, *p* < 10^−10^, Kruskal-Wallis and Dunn-Sidak multiple comparisons test) (Supplementary Fig 1). To test whether this mechanical anisotropy of the endoneurium is due to the longitudinal orientation of morphological structures (e.g., axons), we performed acute crushes on dissected nerve samples in ice-cold solution two hours prior to measurement, to damage those structures while minimising inflammatory responses. Acutely crushing nerves significantly decreased the stiffness of longitudinally measured samples (median *K* = 66 Pa, *p* < 10^−10^, Kruskal-Wallis and Dunn-Sidak multiple comparisons test, relative to naïve longitudinal) but not the stiffness of cross-sectional specimens (median *K* = 56 Pa, *p =* 0.95) (Supplementary Fig 1), suggesting that the structural integrity of axons is a major contributor to the mechanical anisotropy of nerve endoneurium.

### Nerve crush injury leads to a long-lasting increase in nerve stiffness

Crush injuries, or axonotmesis, are one of the two major types of injury a nerve may undergo^24,25^. Compression of the nerve causes damage to the tissue, disrupting axon integrity and resulting in long-term loss of nerve conduction. Crush injury triggers an inflammatory response, leading to changes in the cellular and ECM composition at the injury site. This response is a vital step leading to axon regeneration and restoration of peripheral nerve function. To test whether these injury-associated changes in tissue composition may be associated with changes in tissue mechanics, we carried out a compression injury in the sciatic nerve of rats and evaluated accompanying changes in tissue stiffness and ECM composition at the crush site 3 days as well as 90 days post injury.

The injury immediately eliminated all sciatic nerve-related motor control and sensory perception in the operated limb. Parameters were assessed 3 days post-injury, as by that time point inflammation, Schwann cell transdifferentiation into Bungner cells, and removal of axonal debris are already underway, but new axons have not yet extended through the crush site ^19,20^. We also studied nerves at 90 days post-injury, by which time regeneration had already successfully occurred and animals had recovered sensory and motor control over the operated limb.

While the stiffness of acutely crushed nerves was similar to that of controls (Supplementary Fig 1), at 3 days post-crush, endoneurial stiffness had significantly increased from *K* = 66 Pa to 138 Pa (*p* < 10^−10^, Kruskal-Wallis and Dunn-Sidak multiple comparisons test, relative to naïve nerve) (Fig 2A-B), indicating that changes in tissue stiffness were not directly caused by the damage to the nerve architecture but rather by inflammatory and tissue remodelling processes (Fig 2A).

**Fig 2.**
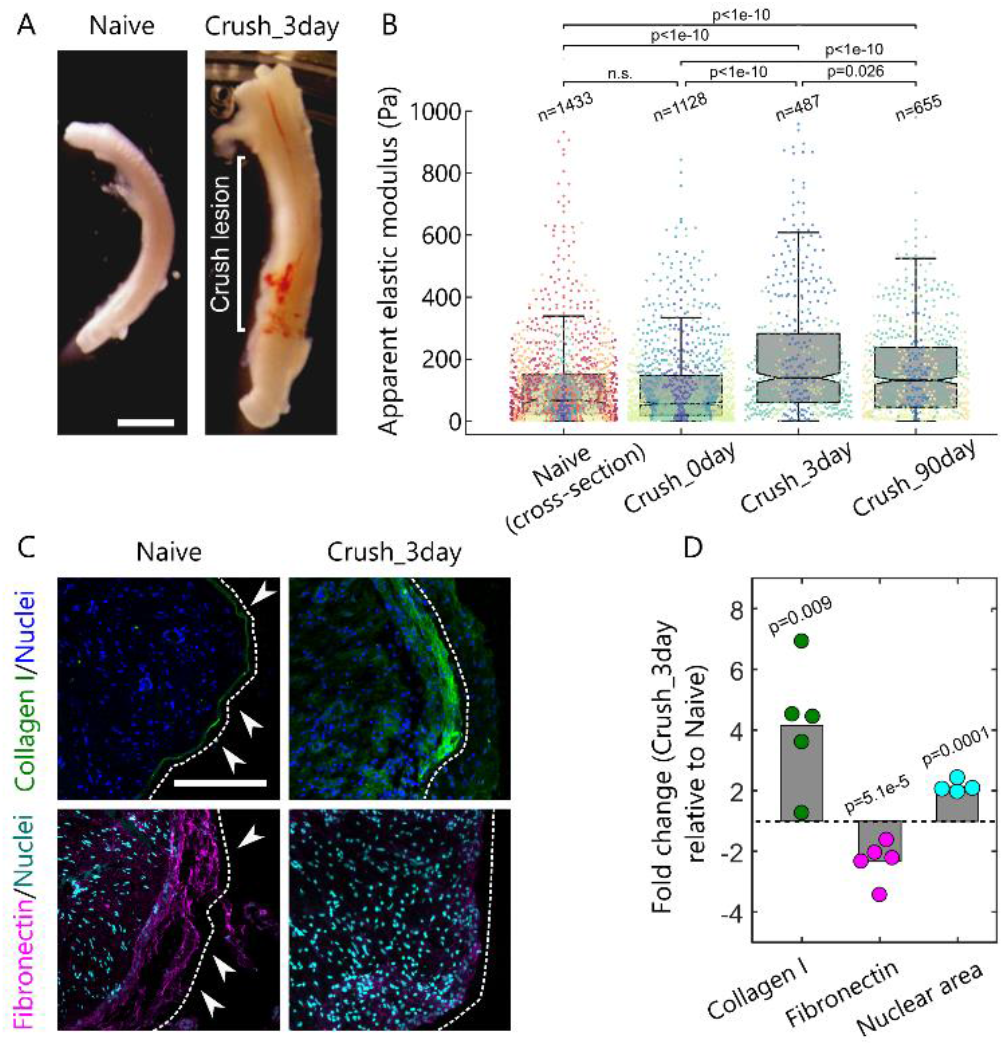
Crush injury leads to long-term stiffening of peripheral nerve tissue. (A) Sciatic nerve 3 days after crush injury and a naïve control. Swelling of tissue is visible in the injured nerve. Scale bar: 2 mm. (B) Tissue stiffness for naïve nerves, acutely crushed nerves, 3 days post-crush injury, and 90 days post-crush injury. Nerve tissue significantly stiffened after injury (*p* < 0.001, Kruskal-Wallis and Dunn-Sidak multiple comparisons test); mechanical changes lasted until chronic timepoints. Measurements *n* represented by individual dots; measurements taken from the same animal are depicted with the same colour (naïve En replotted from Fig. 1F, Crush_0day *N*=4 animals, *n*=1128 measurements, Crush_3day *N*=3, *n*=487, Crush_90day *N*=6, *n*=655). n.s.: not significant (p values > 0.05). (C) Cross-sections of naïve nerve and crushed nerve 3 days post injury. Tissue stained for the ECM components collagen I (green) and fibronectin (magenta), as well as for cell nuclei (Hoechst, blue/cyan). ECM in naïve nerves was largely restricted to the epineurium surrounding the nerve (white arrowheads) but was found throughout the cross-section following injury. Scale bar: 200 μm. (D) Relative changes in ECM (collagen I and fibronectin; *N* = 5) and nuclear (*N* = 4) densities 3 days post crush injury. Collagen levels and nuclear densities significantly increased after crush-injury (*p* < 0.01 and <0.001, respectively, two-tailed Student’s t-test), while fibronectin levels decreased (*p* < 0.001). Bars indicate mean values, data points represent values for individual animals.

To investigate which morphological changes accompanied the observed changes in tissue mechanics, we next investigated selected cellular and ECM components using immunohistochemistry (Fig 2C-D), focusing on collagen I, which is a major contributor to tissue stiffness in many tissues^26^, and on nuclear density, which has been previously reported to contribute to higher tissue stiffness in spinal cord^18^ and developing brain tissue^8,9^. Additionally, we characterised changes in fibronectin – an ECM protein abundantly present in nerves before and after injury^27^ but with unknown contribution to nerve stiffness. We found a significant increase in collagen, which now extended throughout the nerve cross section I (*p* = 0.009, Student’s t-test) and decrease in fibronectin (*p* = 5.1×10^−5^) compared to naïve nerve controls. Furthermore, the relative area occupied by cell nuclei increased significantly 3 days post injury (p = 0.0001) (Fig 2C-D).

90 days post injury, when limb function had fully recovered, nerve tissue exhibited a small but statistically significant decrease in tissue stiffness relative to the 3 days timepoint (131 Pa compared to 138 Pa, *p =* 0.026, Dunn-Sidak multiple comparisons test; Fig 2B) but remained stiffer than naïve tissue (*p* < 10^−10^), indicating that crush injury-related tissue stiffening does not fully disappear upon nerve regeneration.

### Transection injury leads to a large increase in nerve stiffness

Having identified a stiffening of nerves in response to crush injury, we assessed whether this pattern was also seen in another major type of nerve injury. Transection injury, also known as neurotmesis, occurs when a nerve becomes fully severed, resulting in the formation of two nerve stumps with a gap between them. The subsequent healing response by the body requires the formation of a tissue bridge, formed by cells migrating from the nerve stumps and eventually becomes the substrate within which axons regrow across the lesion. During the regenerative response, the nerve tissue bridge is a distinct tissue type with unique cell and ECM content; it becomes similar in composition to nerve tissue over a time period of months^28^.

To test how tissue mechanics changes after transection injuries, we generated a 5-mm gap between the stumps of transected rat sciatic nerves *in vivo* and implanted a PDMS conduit to house the nerve bridge and guide nerve regeneration (Fig 3A). Measurements of both nerve bridges and stumps were done at 5 days post-transection, at which time point stumps are still distinguishable and bridges contain macrophages, fibroblasts, a dense ECM, and Bungner/Schwann cells migrating from the nerve stumps, while regenerating axons are not yet found^19,28^. We also evaluated bridges at 90 days post-surgery, when regeneration is complete, and animals had recovered limb sensation and motion.

**Fig 3.**
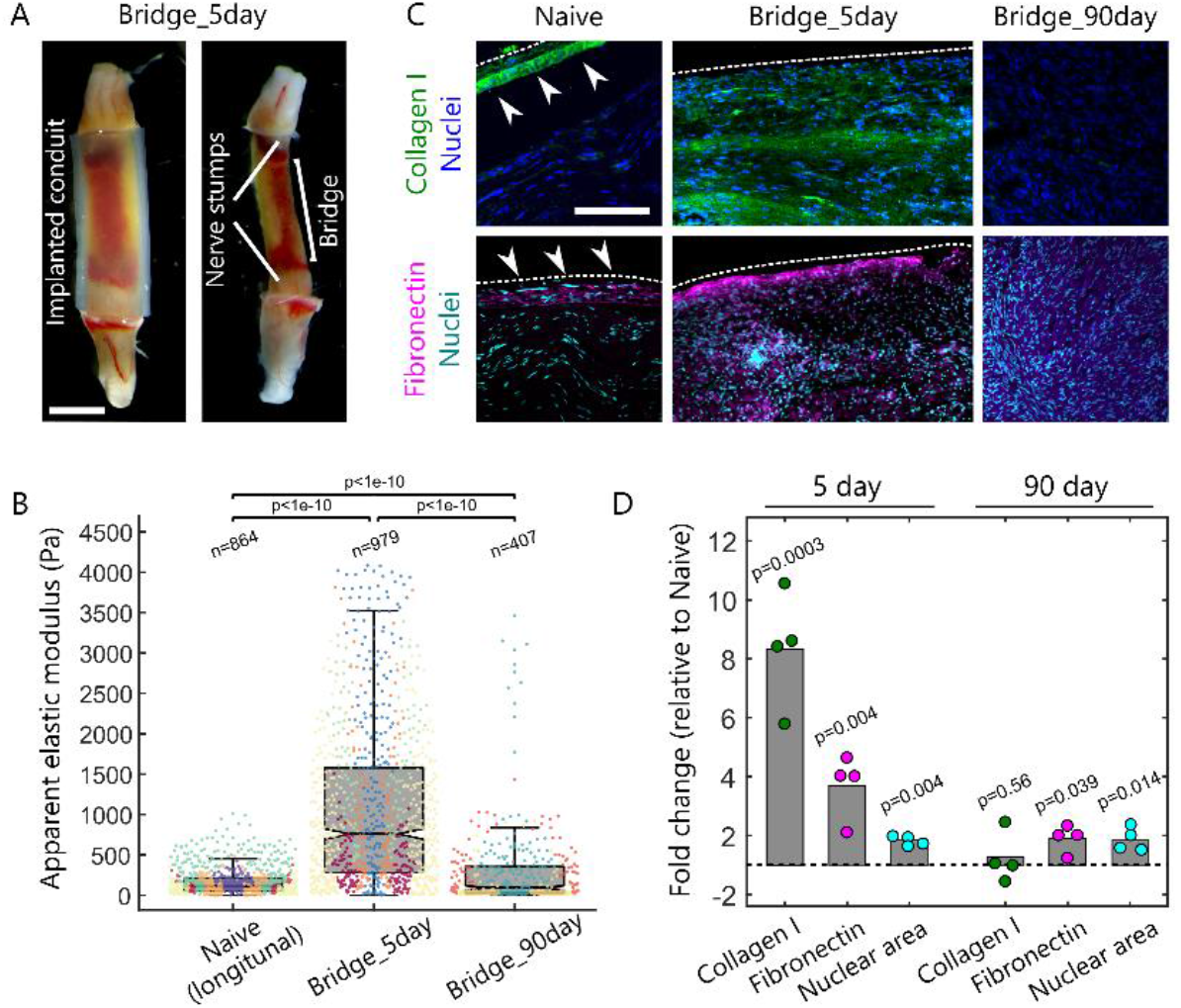
Transection injury leads to the formation of a stiff nerve bridge. (A) Sciatic nerve 5 days following transection injury. The tissue bridge connecting the two nerve stumps is visible after removal of the silicone conduit (right). Scale bar: 2 mm. (B) Tissue stiffness for transection injuries. Nerve bridges formed after transection injuries were significantly stiffer than healthy tissue 5 days post injury (*p* < 0.001, Kruskal-Wallis and Dunn-Sidak multiple comparisons test); tissue stiffness decreased towards that of healthy nerve at chronic timepoints. Measurements *n* represented by individual dots, with measurements taken from the same animals depicted with the same colour (naive *N*=8 animals, *n*=864 measurements, Bridge_5day *N*=5, *n*=979, Bridge_90day *N*=4, *n*=407). (C) Longitudinal sections of naïve nerves and transection-injured nerves 5 and 90 days post injury. Tissue stained for the ECM components collagen I (green) and fibronectin (magenta), as well as cell nuclei (Hoechst, blue/cyan). ECM in naïve nerves was largely restricted to the epineurium surrounding the nerve (white arrowheads) but was found within the nerve 5 and 90 days following injury. Scale bar: 200 μm. (D) Changes in ECM (collagen I and fibronectin) and nuclear densities (*N* = 4) at 5 and 90 days post injury relative to healthy controls. Collagen and fibronectin levels as well as nuclear densities significantly increased 5 days after transection injury (all *p* < 0.01, two-tailed Student’s t-tests). Levels went back towards those of the control at 90 days post injury, although fibronectin levels and cell densities were still significantly increased (*p* < 0.05). Bars indicate mean values, data points represent values for individual animals.

As after crush injuries, we found that nerve bridges were significantly stiffer than naïve control nerves (median *K* = 759 Pa) 5 days post-injury (*p* < 10^−10^, Kruskal-Wallis and Dunn-Sidak multiple comparisons test) (Fig 3B). The stiffness of these nerve bridges was even considerably higher than that of regenerating, stiffened tissue of crushed nerves (Figs. 2B and 3B). Also after transection injuries, the observed greater stiffness in bridges was accompanied by changes in ECM composition (Fig 3C, D). Nerve bridges exhibited significantly geater collagen I (*p* = 0.0003, Student’s t-test) and nuclear densities than control nerves (*p* = 0.011). Here, we also observed an increase in fibronectin fluorescence (*p* = 0.004, Student’s t-test relative to naïve samples) (Fig 3D).

After 90 days of recovery, the stiffness of nerve bridges was significantly lower than at 5 days post injury (*K* = 93 Pa compared to 759 Pa; *p* < 10^−10^, Kruskal-Wallis and Dunn-Sidak multiple comparisons test), dropping to similar (but still significantly higher) stiffness values compared to healthy tissue (*K* = 66 Pa *p* < 1×10^−10^, Kruskal-Wallis and Dunn-Sidak multiple comparisons test) (Fig 3B). In line with this observation, collagen I levels decreased to levels similar to those of naïve nerve tissue (*p* > 0.05relative to naïve tissue, two-tailed Student’s t-test) (Fig 3D). The relative area occupied by nuclei, however, remained similar to that found in the bridge tissue at 5 days post-transection and higher than in the healthy control (*p* = 0.014 relative to naïve, two-tailed Student’s t-test). Fibronectin values also decreased but remained significantly higher than naïve nerve levels (*p* = 0.039, two-tailed Student’s t-test) (Fig 3D).

Nerve transection injuries also cause an inflammatory response at the nerve stumps. Damage-associated changes in the stumps are important to regeneration following a transection injury, as macrophages, fibroblasts and Schwann cells become activated and migrate into the gap from the stumps to form the nerve bridge^28^. We evaluated the stiffness of the nerve stumps at 5 days post transection, after the nerve bridge had already formed. Tissue stiffness exhibited a well-defined boundary at the stump-to-bridge border, with stumps exhibiting a significantly lower stiffness than nerve bridges with a median *K* = 202 Pa (*p* < 10^−10^, Kruskal-Wallis and Dunn-Sidak multiple comparisons test) (Supplementary Fig 2). However, stump stiffness was significantly higher than that of healthy nerve tissue (*p* < 10^−10^, Mann-Whitney U-test) and closer in value to nerve crush injuries (*p* = 1.8×10^−4^, Kruskal-Wallis and Dunn-Sidak multiple comparisons test), suggesting that similar cellular responses might be at play in both injury models, leading to similar mechanical changes of nerve tissue.

### Schwann cells polarize on substrates mechanically mimicking injured tissue

Schwann cells are one of the key contributors to nerve regeneration in the PNS. Upon injury, they transdifferentiate from an axon-supporting cell type into mediators of nerve regeneration. In this state – often referred to as Bungner cells^20^ – Schwann cells polarize, extending processes and becoming proliferative and motile to guide and remyelinate regenerating axons across the injured nerve. As our results showed that injured nerves experienced significant increases in stiffness (Figs. 2-3), we tested whether a stiffened environment may serve as a cue to trigger Schwann cell transdifferentiation.

Schwann cells isolated from primary tissues are usually expanded in tissue culture flasks prior to seeding them into Petri dishes. However, tissue culture flasks have elastic moduli in the GPa range and are thus ∼10^7^ orders of magnitude stiffer than nerve tissue. This mechanical mismatch may potentially prime Schwann cells and alter their mechanosensitive response to the stiffness of the substrates they are subsequently cultured on. To avoid this, we here developed a protocol which allowed us to culture acutely isolated primary Schwann cells directly onto soft polyacrylamide substrates of varying stiffnesses for 6 days immediately after extraction from their host tissue before we evaluated changes in morphology, proliferation, and migration.

On the softest substrates, which were similarly soft as naïve nerve tissue (∼0.1 kPa), Schwann cells remained spherical, a morphology typically found in Schwann cells cultured in differentiation-inhibiting media^29^. When substrate stiffness was as high as in post-injury nerve tissue (∼1 kPa), however, Schwann cells polarized and extended long processes, a morphological characteristic typically associated with Bungner cells^30^ (Fig 4A-D). Process lengths significantly increased further at even higher substrate stiffnesses (*p* < 10^−10^, one-way ANOVA) (Fig 4E). Also, focal adhesion area, a measure of cell adhesion, and cell proliferation both increased with increasing substrate stiffness (*p* = 5.1×10^−5^, one-way ANOVA) (Fig 4F), as seen during the transition of Schwann cells to Bungner cells in response to injury^20,30^.

**Fig 4.**
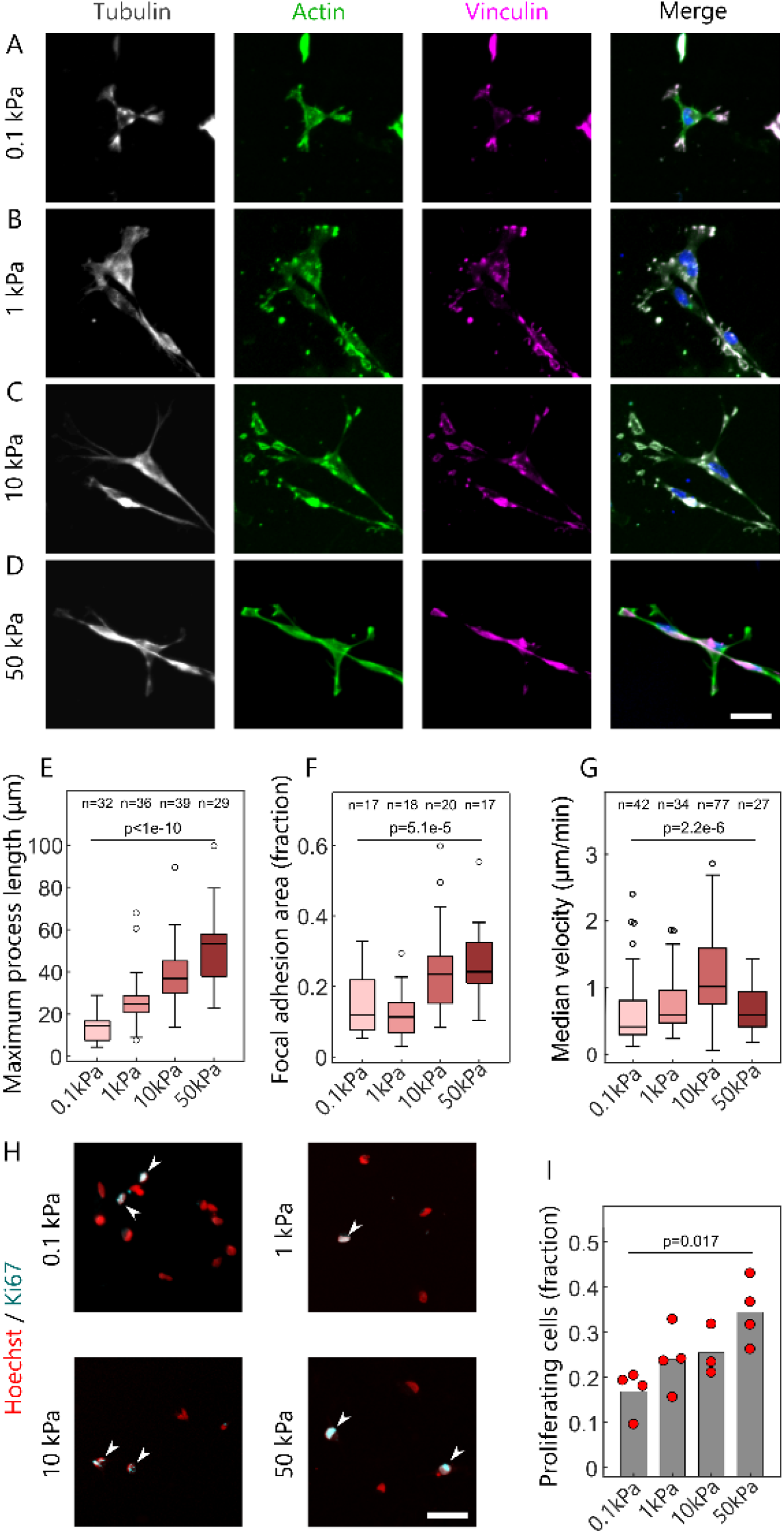
Environmental stiffness regulates primary Schwann cell behaviour. (A-D) Schwann cells cultured on polyacrylamide substrates of (A) 0.1, (B) 1, (C) 10, and (D) 50 kPa shear modulus at 6 days *in vitro*. Cells were stained for cytoskeletal components (actin, tubulin) and focal adhesions (vinculin). Scale bar: 25 μm. (E-G) Quantification of (E) maximum Schwann cell process lengths and (F) focal adhesion areas at 6 days *in vitro* as well as of (G) Schwann cell motility at 1 day *in vitro. n* = number of cells from 3 – 5 biological replicates. (H) Schwann cells on substrates of different stiffnesses stained for nuclei (Hoechst) and proliferation markers (Ki67, arrow heads). Scale bar: 50 μm. (I) Quantification of cell proliferation (dots represent average values for different cultures). (E-G, I) All parameters increased significantly with increasing substrate stiffness (one-way ANOVA), with cell motility reaching a peak at 10 kPa substrate stiffness, suggesting that Schwann cells polarized and became more proliferative and motile on substrates with a stiffnesses exceeding that of healthy nerve tissue.

Live imaging of GFP+ primary Schwann cells revealed that also cell migration significantly depended on substrate stiffness (*p* = 2.2×10^−6^, one-way ANOVA), reaching its peak at 10 kPa (Fig 4G). Finally, cell proliferation, which is increased in Bungner cells, also increased with substrate stiffness (*p* = 0.017, one-way ANOVA) (Fig 4H-I). These results suggested that the mechanical changes seen in injured nerves may help regulating the transition from static Schwann cells to highly motile Bungner cells, thus facilitating regeneration.

## Discussion

In this study, we characterised mechanical changes of peripheral nerve tissue before and after crush and transection injuries using AFM-based indentation experiments. We found that, in healthy nerves, endoneurium, perineurium, and interfascicular epineurium were similarly soft with an elastic modulus of *∼* 60 Pa. With such a low modulus, peripheral nerves belong to the softest vertebrate tissues^31,32^, and they are similarly soft as brain^17^ and spinal cord^6^ white matter, which is also mainly composed of myelinated axons. Only the epifascicular epineurium had a higher stiffness, almost double that of the other nerve compartments (∼100 Pa) (Fig. 1). After both crush injuries and transection injuries (Figs. 2-3), the most common and damaging types of injuries seen in the clinic^33^, nerve tissue stiffened significantly within a few days after injury.

This increase in tissue stiffness was accompanied by changes in cellular densities and ECM composition. In healthy nerve tissue, the endoneurium is composed primarily of axons and Schwann cells, with only a sparse collagen and proteoglycan ECM^34^, and the perineurium is mainly made of layers of perineurial cells^35^. Both epi- and interfascicular epineuria, on the other hand, are primarily composed of collagen I-rich ECM. The difference in mechanical properties between the epineuria can likely be explained by the structure of their ECM: aligned bundles of collagen I in the epifascicular epineuria, able to resist tension along the nerve^20,33,35,36^, will be stiffer than the sponge-like structure of the ECM in the interfascicular epineuria^36^, able to absorb compressive forces^21,37^.

Upon injury, cell densities and collagen I content of the damaged nerve tissue increased. Collagen I is a structural ECM component, which is abundantly found in most scar tissues^38^ (except in scars of the CNS^5^) and whose abundance strongly correlates with stiffness in many tissues^26^. Cell densities have been reported to scale with the stiffness of neural tissues^8,9,18,39^, although negative correlations between cell body densities and CNS tissue stiffness have also recently been reported^40^. In addition to the composition of a tissue, other factors, such as the structure and arrangement of cellular and extracellular tissue components might be crucial determinants of tissue mechanics.

Tissue stiffness modulates cell function *in vivo*, influencing, amongst others, cellular motility and activity^8,10,14^. Previous studies investigating the response of Schwann cells to substrate stiffness found complex, non-trivial, and sometimes inconsistent adaptations of cell morphology, adhesion, and motility to their mechanical environment^41–44^. However, in these studies, cells were usually exposed to tissue culture plastics (∼GPa substrate stiffness) during the expansion phase, and the softest substrates used had elastic stiffnesses in the kPa range, mechanically similar to or even stiffer than post injury nerve tissue (Figs 2, 3). We here avoided the use of tissue culture plastics and exclusively used substrates whose stiffness was matched to the stiffness of healthy and post injury nerves identified in this study.

Primary Schwann cells polarized and became more motile and proliferative on substrates with a stiffnesses above that of healthy nerve tissue (Fig 4), suggesting that the observed increase in nerve tissue stiffness after injury could contribute to activating Schwann cells, thus facilitating nerve regeneration. Focal adhesions are known mediators of Schwann cell activity in development and regeneration via molecules such as Focal Adhesion Kinase and downstream effectors such as the mechanosensitive transcriptional regulators YAP/TAZ^45,46^. We found a higher abundance of focal adhesions in Schwann cells on substrates with stiffnesses close to that of injured nerves (Fig. 4F), suggesting a possible molecular mechanism contributing to their regenerative response.

While the baseline stiffness of CNS white matter^5,11^ and peripheral nerves is similar (Fig 1), injury leads to opposite mechanical responses of the tissues. Adult mammalian PNS tissue becomes stiffer after injury (Figs 2-3), whereas adult mammalian CNS tissue becomes softer^5,6^. An increased stiffness of peripheral nerves after injury may not only lead to enhanced Schwann cell proliferation, motility, and polarization (Fig 4), thus extrinsically supporting neuronal regeneration, but also to enhanced axon growth^8^, while the soft environment mammalian neurons encounter after injuries to the CNS may slow down axon growth and limit the promotion of pro-regenerative factors released by glial cells. In line with this thought, injuries to the spinal cord of zebrafish, which in contrast to the mammalian spinal cord can regenerate after injury, lead to the stiffening of the tissue^7^. Our results thus suggest that, in addition to chemical signals^4^, the increase in nerve tissue stiffness after injury may significantly contribute to facilitating neuronal regeneration, and that future approaches to treat spinal cord injuries should include strategies to stiffen the injury site.

## Materials and Methods

### Surgical procedures and sciatic nerve tissue harvesting

All animal procedures carried out were in compliance with the United Kingdom Animals (Scientific Procedure) Act of 1986 and institutional guidelines. Sciatic nerves were harvested from ∼250g female rats. Lewis strain rats (Charles River UK) were used when surgical procedures were carried out prior to tissue harvesting, while Sprague Dawley rats (Charles River UK) were used when naïve tissue was used.

Animals undergoing surgical procedures were housed in groups of 5 and provided *ad libitum* access to water and food for at least 7 days prior to surgery. Anaesthesia was induced and maintained using isoflurane in oxygen delivered via a facemask, at the beginning of which animals received a subcutaneous injection of the non-steroidal anti-inflammatory drug meloxicam (1.5 mg/ml). Body temperature was monitored and maintained at 37ºC during anaesthesia using a rectal probe and thermal blanket. During surgery, the right sciatic nerve of animals was accessed dorsally around the site of the nerve trifurcation, accessed through the septum between the biceps femoris and vastus lateralis muscles, and either a nerve crush or nerve gap injury was carried out for each animal.

Nerve crush injury was carried out by applying pressure using tube occluding forceps equipped with a ratchet mechanism to ensure constant application of force during and between crushes. Tissue was crushed for 10 seconds starting at 2 mm proximal to the trifurcation. A total of four sequential crushes were carried out proximal to that point until the lesion reached a total length of 7 mm. In sciatic nerve gap injuries, the sciatic nerve was cleanly transected using surgical scissors 2 mm and 7 mm proximal of the trifurcation. The resulting nerve stumps were placed inside a 6 mm-long silicone tube (2 mm diameter, Portex), and sutured ∼0.5 mm into the tube (three or two sutures through tube and nerve epineurium, 9/0 nylon Ethicon), resulting in a 5 mm-long nerve gap with no additional nerve tension or slack introduced. All animals were allowed to recover following surgery, and provided a further dose of meloxicam the day after surgery. Animals were sacrificed after a post-operative period of 3 days or 90 days (nerve crushes), or 5 days or 90 days (nerve gap injuries).

At the end of the post-operative recovery period (or immediately, for naïve animal tissue) animals were sacrificed and the tissue was collected for atomic force microscopy and/or stain intensity analysis (immunohistochemistry). Animals were sacrificed by overdose of euthatal (pentobarbitone) administered intraperitoneally, followed by neck dislocation. Euthatal was combined at a 1:1 v/v ratio with lidocaine anaelgesic and delivered at a total dose of 3 ml/kg of bodyweight. Anaesthetic overdose was chosen as a method of sacrifice instead of rising CO_2_ concentration, as the latter has been associated with acidosis and tissue stiffening^47^. Sciatic nerves (approximately a 1 cm fragment proximal to the trifurcation) were immediately dissected. The fragment was split into two, with one half being immediately fixed in paraformaldehyde and processed for immunostaining (see below), and the other half being transferred into mammalian buffer saline (for AFM, see below) on ice. In the case of nerve injury tissue, the contralateral sciatic nerve was also collected to use as a naïve control. All AFM measurements were carried out within 5 hours of sacrifice, minimising tissue degradation.

### Atomic force microscopy indentation experiments

Harvested sciatic nerves were transferred into dishes containing ice-cold mammalian physiological saline (121 mM NaCl, 5 mM KCl, 1 mM MgCl_2_, 1 mM CaCl_2_, 0.4 mM NaH_2_PO_4_, 23.8 mM NaHCO_3_, 5.6 mM glucose, previously described by others^48^). Mammalian physiological saline was prepared freshly prior to the experiment. To establish a pH of 7.3, a gas mixture of 95% CO2 and 5% O2 was bubbled through the solution. Under a dissection microscope, external blood vessels and any excess tissue surrounding the nerve were removed, and the nerve stumps were trimmed off with a steel blade.

The remaining nerve was then cut into ∼3 mm fragments for mounting and sectioning. Just prior to cutting into fragments, some nerves (Crush_0day condition, Supplementary Fig 1 and Fig 2) were crushed for 10 seconds using forceps, in a similar fashion to the crush injury protocol described above. Nerve fragments were embedded in warm 4% w/w low melting point agarose (A9539, Sigma) in phosphate buffered saline (PBS). Once cooled and hardened, the agarose was trimmed into blocks and mounted onto a vibrating microtome filled with chilled mammalian physiological saline, where nerve fragments were cut into 500 μm sections, either longitudinally or as cross-sections. Sections were stained for axons for 1 hr at room temperature with fluoromyelin (1:250 v/v in mammalian physiological saline; Invitrogen, F34651). These were then mounted onto 35 mm plastic dishes (Z707651, Sigma) with thin strips of cyanoacrylate glue (Super Glue, Loctite), which adhered to the agarose, and filled with room temperature mammalian physiological saline. Nerve samples used for epifascicular epineurium measurements were not embedded in agarose or sectioned, and simply mounted on their side on a bed of agarose on a plastic dish.

To perform indentation experiments, dishes containing mounted nerves were transferred to an optical inverted microscope (Axio Observer.A1, Carl Zeiss Ltd.) equipped with a motorised xy stage. The sample was imaged with a CCD camera (The Imaging Source GmbH), and a region of interest was defined based on the fluoromyelin stain pattern. A custom python script broke down this area of interest into a grid of 20 × 20 μm squares. At the centre of each of these squares an elasticity measurement was carried out sequentially using a cantilever probe mounted on a JPK Nanowizard Cellhesion 200 AFM (JPK Instruments AG). After indentation measurements, some naïve sciatic nerve samples were fixed and immunostained to identify nerve compartments and confirm the fluoromyelin stain pattern was accurate (see below).

Measurements were taken using tipless cantilever probes, onto which a polystyrene bead (37.28 ± 0.34 µm diameter; Microparticles GmbH) had been previously glued (ultraviolet curing, Loctite). Arrow-TL1 cantilevers were used (NanoSensors, with a spring constant of ∼0.02 N/m), for each of which the exact spring constant was calculated using the thermal noise method^49^. For each elasticity measurement, the cantilever probe was lowered onto the sample at a speed of 10 μm/s, indenting it until a force threshold of 10 nN was reached. The probe was then retracted at a speed of 50 μm/s, and the motorised stage moved the sample to allow for a new measurement to be taken in the neighbouring grid square. The resulting force-distance curves were later processed using a previously described custom MATLAB (Mathworks, R2008a) script^22^. Applying the Hertz model^50^ and modelling the cantilever as a sphere indenting a half space, an elasticity value in the form of apparent reduced elastic modulus *K* was calculated from the force applied *F*.

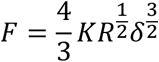

For the Hertz model to be valid, strain applied to the sample must be small and indentation depth must be much smaller than sample thickness. Elasticity was calculated for an indentation depth δof 2 µm, which resulted in a contact radius *a* =√*R*δof approximately 6 µm, with *R* being the radius of the bead. Biological materials tend to begin exhibiting deviations from elastic behaviour at compressive strains ε = 0.2*a*/*R* > 0.02, which places the measurements taken within an acceptable range. The condition *a*/*h* < 0.1, *h* being the thickness of the sample analysed, is also satisfied which is necessary to treat the sample as a half space^51^.

Elastic modulus measurement plotting and statistical analysis was carried out in MATLAB, and presented as boxplots with overlaid scatter plots. Box plots consist of a box containing the 25th to 75th percentile range of measurements (the interquartile range), an inner box mark indicating the median value, and whiskers of length 1.5 times the interquartile range containing most of the remaining data points. In overlaid scatter plots, each dot represents one measurement, with dots with the same colour representing measurements of tissue from the same animal.

### Immunohistochemistry

All tissue was fixed by immersion in paraformaldehyde solution (40 mg/ml in PBS) at 4 ºC overnight, prior to sectioning and staining. Tissue samples were then moved into a sucrose solution (30% w/w in PBS; S0389, Sigma) for cryoprotection for at least 16 hours at 4 ºC, until further processing. Tissue was mounted on optimal cutting temperature compound (Tissue-Tek, 4583), transferred to a cryostat (CM3050 S, Leica), and sectioned into 12 µm sections. Tissue sections were transferred to glass slides, allowed to dry at room temperature overnight, and stored at -20 ºC until used for immunostaining. Tissue samples previously used for AFM were already sectioned using a vibratome as part of the indentation experiments sample processing, and were therefore not cryoprotected or further sectioned. Stain intensity analysis (quantification of ECM components, cell density) was carried out on cryosectioned samples that had not been used for AFM. Immunostaining of samples post-indentation measurements was only used to identify nerve compartments.

In preparation for immunostaining, sections were permeabilised with Triton X-100 solution (Sigma, T8787). Samples were blocked in blocking buffer (tris-buffered saline containing 0.03% v/v Triton X-100 and 10% v/v donkey serum [Millipore, s30-100ml], for 1 hr at room temperature) to minimise non-specific antibody binding. Samples were then incubated in primary antibodies (see Supplementary Table 1) in antibody buffer (10% blocking buffer v/v in PBS) overnight at 4 ºC, covered in paraffin film to prevent drying. Primary antibodies were removed by washing in PBS and Triton X-100 solution, and secondary antibodies in antibody buffer were added for 2 hr at room temperature. After a further two washes in PBS and a single wash in non-saline Tris-buffered solution (T6066, Sigma), samples were encased with a glass coverslip and Fluorsave mounting agent (Millipore, 345789), and stored at 4 ºC until imaged. Unless otherwise specified all washes were performed 3 times, for 10 minutes each.

Imaging was carried out using a confocal microscope (Leica TCS SP5). Fluorescence images were taken using a camera and exported for analysis into Image-J software package (v1.48, National Institutes of Health, USA). Cell density was measured using an automated custom macro in Image-J, applied over a 100 × 100 µm area, and presented as relative nuclear density (fraction of the image covered by cell nuclei). Three measurements were averaged to produce an *n* = 1 biological replicate corresponding to one animal. Stain intensity analysis was carried out by selecting a 100 × 100 µm area within the tissue, and subtracting a background intensity measurement (taken from a region of the image not containing any tissue). Measurements from injured nerves were compared to those of naïve nerves and a fold change in stain intensity was calculated. Data was analysed and plotted in MATLAB (Mathworks, R2016b). For post-AFM samples, images of the nerve samples were exported, compared, and aligned to those taken using the indentation experiment setup. This allowed for stiffness measurement maps to be aligned to immunostained sections, and was used to confirm that the fluoromyelin stain pattern used in the *ex vivo* AFM measurements could accurately identify the nerve compartments.

### Polyacrylamide cell culture substrates

Polyacrylamide substrates on glass coverslips were prepared as previously described^12^ from a combination of acrylamide (40% w/w; A4058, Sigma), bisacrylamide (2%; BP1404-250), and hydroxyacrylamide (97%; 697931, Sigma) solutions, at a ratio of 177:100:23 in PBS (D8537, Sigma). The volume of PBS added to the polyacrylamide mix was controlled to produce a gel of a specific stiffness while having a minimal effect in the chemical properties of the gels across conditions (10.6% polyacrylamide in PBS v/v for 0.1 kPa, 15% for 1 kPa, 30% for 10 kPa. 100% for 50 kPa).

Gel substrates were prepared on 19 mm diameter glass coverslips. These were cleaned by alternate dipping in ddH_2_O and EtOH, treated with 0.1N NaOH (VWR, E584) for 5 min, and functionalised with 150 μl of APTMS (3-Aminopropyltrimethoxysilane; Sigma, 281778) solution for 2.5 min, finished by thorough rinsing in water. Finally, coverslips were treated with glutaraldehyde 0.5% (v/v) solution (G6257, Sigma) in ddH_2_O (15230089, Fisher Scientific) for 30 min at room temperature.

Polyacrylamide gel mixes were then prepared into 500 μl mixes as described above, and degassed under vacuum for 10 min. Polyacrylamide polymerisation was initiated by addition of 5 μl of ammonium persulfate solution (0.1 g/ml in ddH_2_O; Sigma, 281778215589) and 1.5 μl of TEMED (N,N,N’,N’-tetramethylethylenediamine; 15524-010, Invitrogen). A 8 μl drop was quickly deposited onto each 19 mm coverslip, and flattened under a 22 mm diameter glass coverslip (previously treated with RainX [Rain Repellent Krako Car Care International Ltd, UK] to render hydrophobic).

Gels were allowed to swell in PBS overnight, before removing the 22 mm coverslip. Gels were then sterilised for 1 hour under UV light, functionalised with PDL (Poly-D-lysine; 100 μg/ml in PBS; P6407, Sigma) overnight and laminin (1 μg/ml in PBS, Sigma) for 2 hours at room temperature. Gel substrates were used within 2 days of preparation, and allowed to swell in cell culture medium for 30 min prior to plating of any cells.

One substrate of each stiffness for each prepared batch was measured by AFM to confirm their stiffness was the expected value. Stiffness values were calculated as described in the AFM section, using tipless silicon cantilevers, with a spring constant appropriate for the stiffness of each substrate. For 0.1 kPa gels Arrow-TL1 (spring constant of ∼0.02 N/m, NanoSensors), for 1 kPa substrates SICON-TL-20 (spring constant ∼0.29 N/m, AppNano), for 10 and 50 kPa substrates TL-FM-10 (spring constant ∼2.8 N/m, Nanosensors). Arrow-TL1 cantilevers had ∼37 μm beads glued, while SICON-TL-20 and TL-FM-10 had ∼20 μm beads glued (both Microparticles GmbH). Cantilever stiffness and bead diameter values were chosen to ensure the 10 nN force threshold was reached at a depth δ< 5 μm. Stiffness values calculated for polyacrylamide substrates were converted from apparent reduced elastic modulus *K* to shear modulus *G* with the following equations^22^, setting the Poisson ratio *ν* of polyacrylamide to 0.4867^52^.

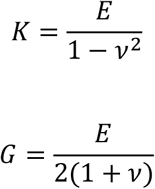

### Schwann cell cultures

Cultures were prepared from postnatal day 1 to 5 Sprague Dawley rat (Charles River UK) sciatic nerves. For Schwann cell migration experiments, green fluorescent protein (GFP) transgenic Sprague Dawley rats were used. Cultures were prepared using a variation of the procedure previously described^53^. All animal procedures carried out were in compliance with the United Kingdom Animals (Scientific Procedure) Act of 1986 and institutional guidelines.

Both sciatic nerves of 10 – 20 animals were dissected using sterilised microscissors. The nerve tissue was dissociated by incubation in collagenase solution (30 min at 37ºC, 2 mg/ml, C9407, Sigma), after which equal parts by volume trypsin solution was added (a further 20 min at 37ºC, T0303, Sigma). An equal volume of deoxyribonuclease was finally added (a further 2 min at 37ºC, D5025, Sigma), after which the dissociated pellet of cells was centrifuged (4 min, 1000 rpm). The supernatant solution was removed and substituted for triturating solution, containing 10 mg/ml bovine serum albumin (A7906, Sigma), 0.5 mg/ml trypsin inhibitor (10109886001, Roche), and 0.02 mg/ml deoxyribonuclease; in which the pellet was re-suspended.

To remove the population of nerve fibroblasts from the cell suspension, cells were again centrifuged and re-suspended for 5 min at room temperature in Dulbecco’s phosphate-buffered saline (14190-094, Invitrogen) supplemented with 5 mg/ml bovine serum albumin, and 1:10 50nm-magnetic-bead antibody solution against rat/mouse CD90.1 (Thy1.1) (Miltenyi Biotec, 120-094-523). Nerve fibroblasts express the marker CD90.1, while Schwann cells do not^23^, allowing for this population of proliferative cells to be selectively removed. Once magnetically-tagged antibodies were selectively bound, the cells were centrifuged and suspended in chilled Dulbecco’s phosphate-buffered saline / bovine serum albumin and run through a magnetic separator column (MiniMACS Separator; Miltenyi Biotec, 130-042-102). The flow-through fraction, with nerve fibroblasts removed, was collected and re-suspended in DMEM (11320-033, Invitrogen) supplemented with a further 4 mM of glutamine (25030032, Invitrogen), 100 mg/ml foetal calf serum (FCS, Invitrogen) and an antibiotic-antimycotic agent (15240-062, Invitrogen). A subset of the cells were plated and stained for S100 and vimentin markers, to confirm that Schwann cells had been successfully isolated. The cells were finally plated on polyacrylamide substrates of one of four stiffnesses at a density of 10,000 cells/cm^2^.

### Immunocytochemistry

Cells were cultured on polyacrylamide substrates of specific stiffness for 6 days, with the medium being changed every two days. At the end of this period, cells were fixed in warm paraformaldehyde solution (40 mg/ml in PBS) for 15 min, and then washed in PBS (3 times, 10 min). Cells were incubated for 30 min in blocking solution (0.03% v/v Triton X-100 [T8787, Sigma], 3% v/v bovine serum albumin [A9418, Sigma], in PBS) to improve antibody binding specificity, followed by primary antibodies (Supplementary Table 1, in blocking solution) overnight at 4ºC. Primary antibodies were washed off (PBS, 3 times, 10 min), and were followed by fluorescently-tagged secondary antibodies (Supplementary Table 1, in blocking solution) for 2 hr at room temperature. After one last set of PBS washes, ended by one wash in non-saline Tris-buffered solution, the cells on substrates were mounted on glass slides with Fluorsave mounting agent (Millipore, 345789). Cells were stored at 4 ºC prior to imaging.

Cell imaging was done using a confocal microscope (Leica TCS SP5, Leica Microsystems, Germany). For each substrate 3 images were taken at random sites with a 20X objective, using DAPI nuclear stain as a guide to ensure cells were present. Gain and exposure settings were maintained throughout all imaging session for each stain.

Quantification of cell stains (cell counting or area selection) was performed in Image-J software package (v1.48, National Institutes of Health, USA) by hand. One image of 50 kPa and one image of 0.1 kPa were opened and for each stain contrast was modified until both achieved a satisfactory stain pattern. This contrast modification was then applied to all images to provide a more accurate and uniform quantification. Data was analysed and plotted in MATLAB (Mathworks, R2016b) box plots or scatter plots. Box plots consist of a box containing the 25th to 75th percentile range of measurements (the interquartile range), an inner box mark indicating the median value, and whiskers of length 1.5 times the interquartile range containing most of the remaining data points. Data points outside of the whiskers (outliers) are marked by circles.

### Schwann cell motility assay

GFP-expressing Schwann cell cultures were transferred to a temperature and CO_2_ controlled chamber mounted on a Leica DMI6000B microscope (Leica Microsystems, Germany) with a motorised stage, at 24 hours post-plating for live imaging. For each substrate, 3 images of green fluorescence were taken at random sites using a 20X objective, before moving to the next substrate. The microscope was programmed to acquire a set of images at the same substrate locations every 5 minutes, for a total of 6 hours, using the Leica Application Suite software (Leica Microsystems).

The resulting movies were imported into Image-J software package, de-shaken using the software’s image stabilizer tool, and underwent cell tracking analysis. Cell tracking of Schwann cells was carried out using the TrackMate (v5.2.0) plugin^54^, set to identify particles of an estimated 15 μm diameter. The median distance covered in each of the resulting cell tracks was quantified, and then analysed and plotted in MATLAB. Tracks shorter than 15 frames were excluded from the analysis.

## Acknowledgements

This work was supported by the UK Medical Research Council (MRC) and the Sackler Foundation (doctoral training grant RG70550), the UK Wellcome Trust (Translational Medicine and Therapeutics PhD Programme Fellowship 109511/Z/15/Z), the European Research Council (Consolidator Award 772426), the UK Biotechnology and Biological Sciences Research Council (Research Grant BB/N006402/1), and the UK Medical Research Council (Career Development Award G1100312/1).

## Author contributions

A.C.L. and K.F. designed the study. A.C. and D.G.B. performed the *in vivo* work. A.C.L. performed AFM measurements and *in vitro* work, and analysed the data. All authors interpreted the data. A.C.L. and K.F. wrote the manuscript, with contribution from all authors.

## Supplementary Information

**Supplementary Figure 1.**
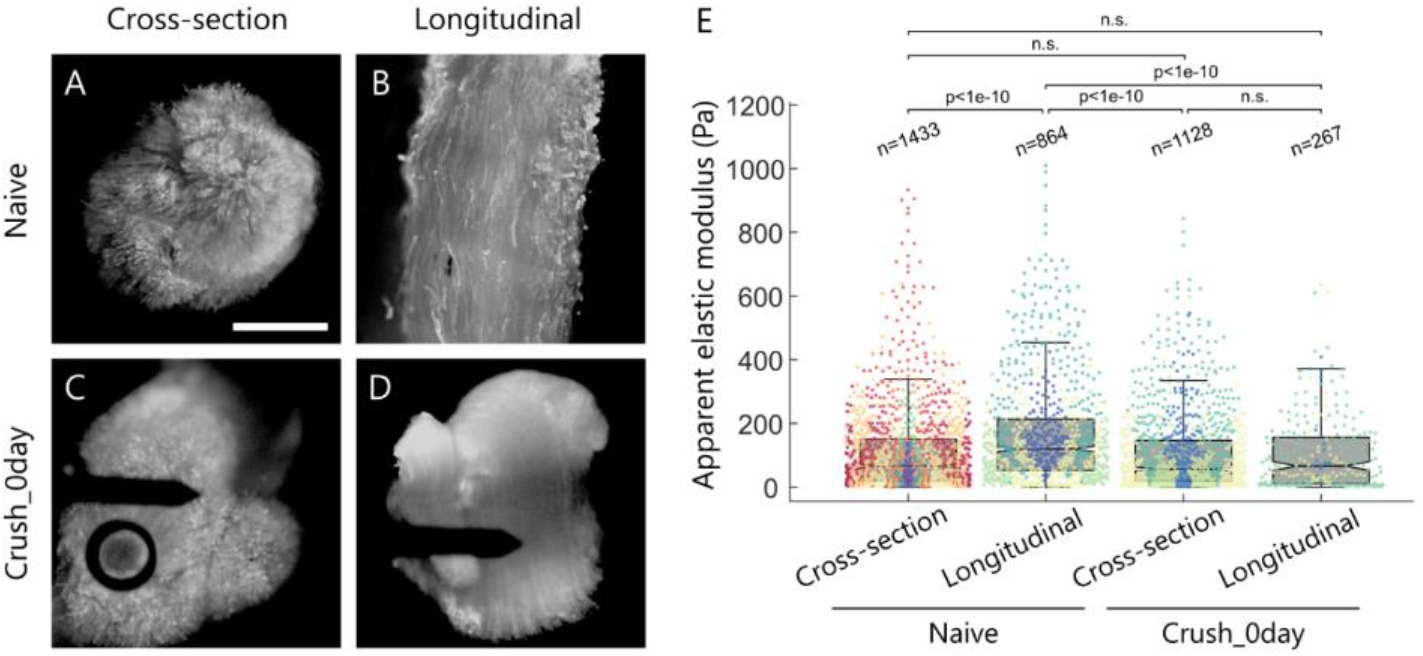
Damage to the nerve structure through acute crushing eliminates the mechanical anisotropy of the endoneurium. (A-D) Live nerve tissue stained with Fluoromyelin; (A, C) cross-sections and (B, D) longitudinal sections. Acute crushes (C-D) disrupt tissue integrity (seen as a loss of detailed staining pattern of individual axons). AFM cantilever visible in (C) and (D). Scale bar: 400 μm. (E) Naïve tissue stiffness was significantly higher when measured on longitudinal endoneurium sections compared to cross-sections (*p* < 0.001, Kruskal-Wallis and Dunn-Sidak multiple comparisons test), indicating mechanical anisotropy. This stiffness difference disappeared in acutely crushed samples (*p* > 0.05), suggesting that axon fibres, which were damaged during the crush, are mainly responsible for the observed mechanical anisotropy of peripheral nerves. Measurements *n* represented by individual dots, with measurements taken from the same animals depicted with the same colours (naive cross-section *N*=16 animals, *n*=1433 measurements, naïve longitudinal *N*=8, *n*=864, Crush_0day cross-section *N*=4, *n*=1128, Crush_0day longitudinal *N*=3, *n*=267).

**Supplementary Figure 2.**
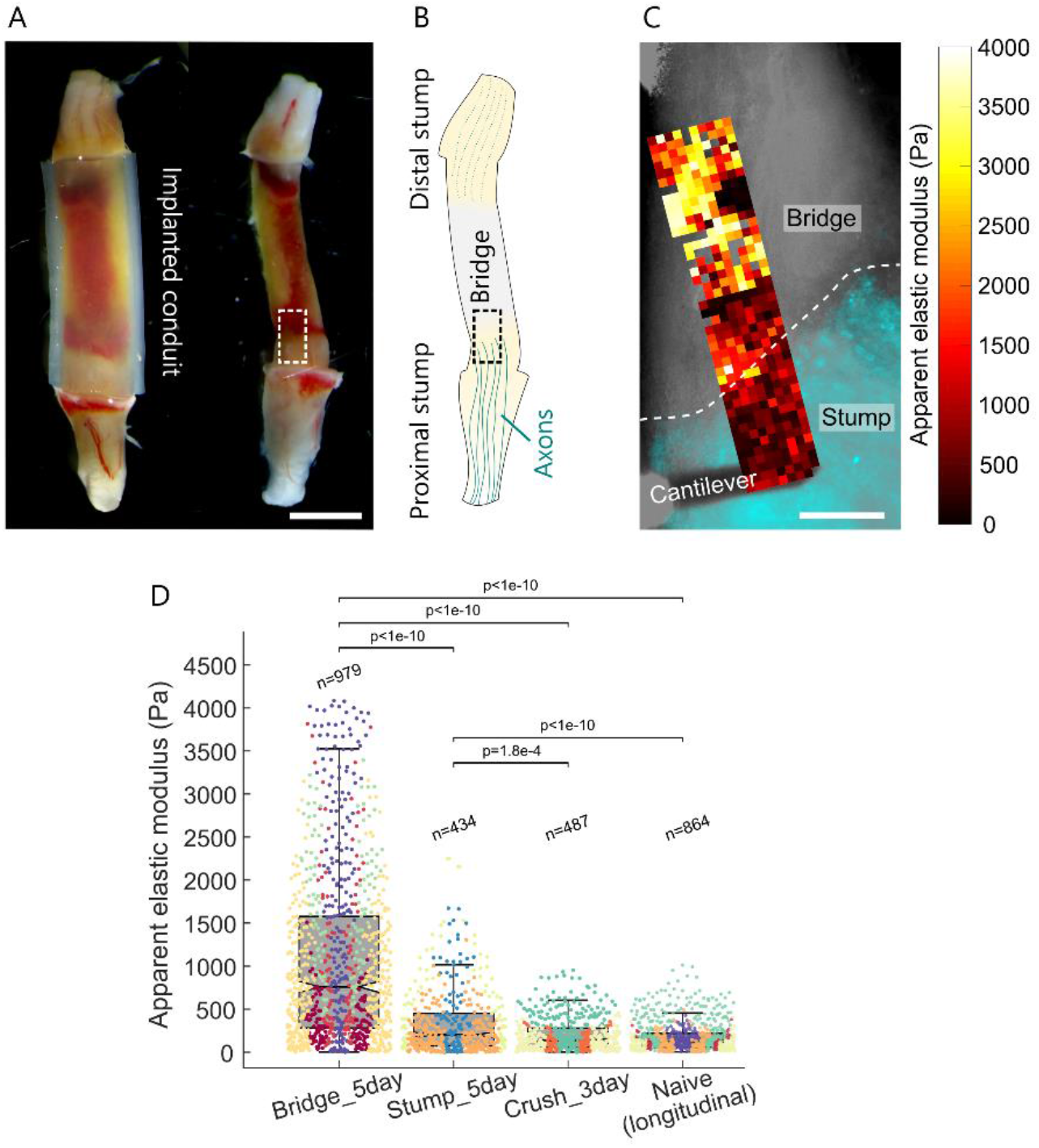
Comparison of nerve tissue mechanics after transection and crush injuries. (A) Images of transected nerve and nerve bridge formed at 5 days post-injury. Scale bar: 2 mm. (B) Schematic of nerve stumps and bridge. Measurements were done over the proximal stump, as axons in the distal stump undergo degeneration at 5 days post-lesion. (C) Longitudinal section of a nerve stump and bridge 5 days after transection injury. The stump was identified by the presence of axons (Fluoromyelin stain; cyan), the nerve stump boundary is indicated by a dashed white line. The AFM Cantilever is visible at the bottom of the image. Overlaid heatmap highlights differences in stiffness between stump and bridge tissue. The region shown here would correspond to that delineated by the dashed squares in (A-B). Scale bar: 200 μm. (D) Tissue stiffness of nerve bridges 5 days post-transection injury, nerve stumps 5 days post-transection injury, and nerve crush tissue 3 days post-crush injury (crush and bridge replotted from Figs. 2B and 3B, respectively; Stump_5day *N*=3 animals, *n*=434 measurements, measurements taken from the same animal depicted with the same colour). With a median *K* = 202 Pa, nerve stumps were mechanically significantly softer than nerve bridge tissue but still significantly stiffer than nerve tissue after crush injuries or naïve tissue (*p* < 0.001 for both, Kruskal-Wallis and Dunn-Sidak multiple comparisons test).

**Supplementary Table 1.**
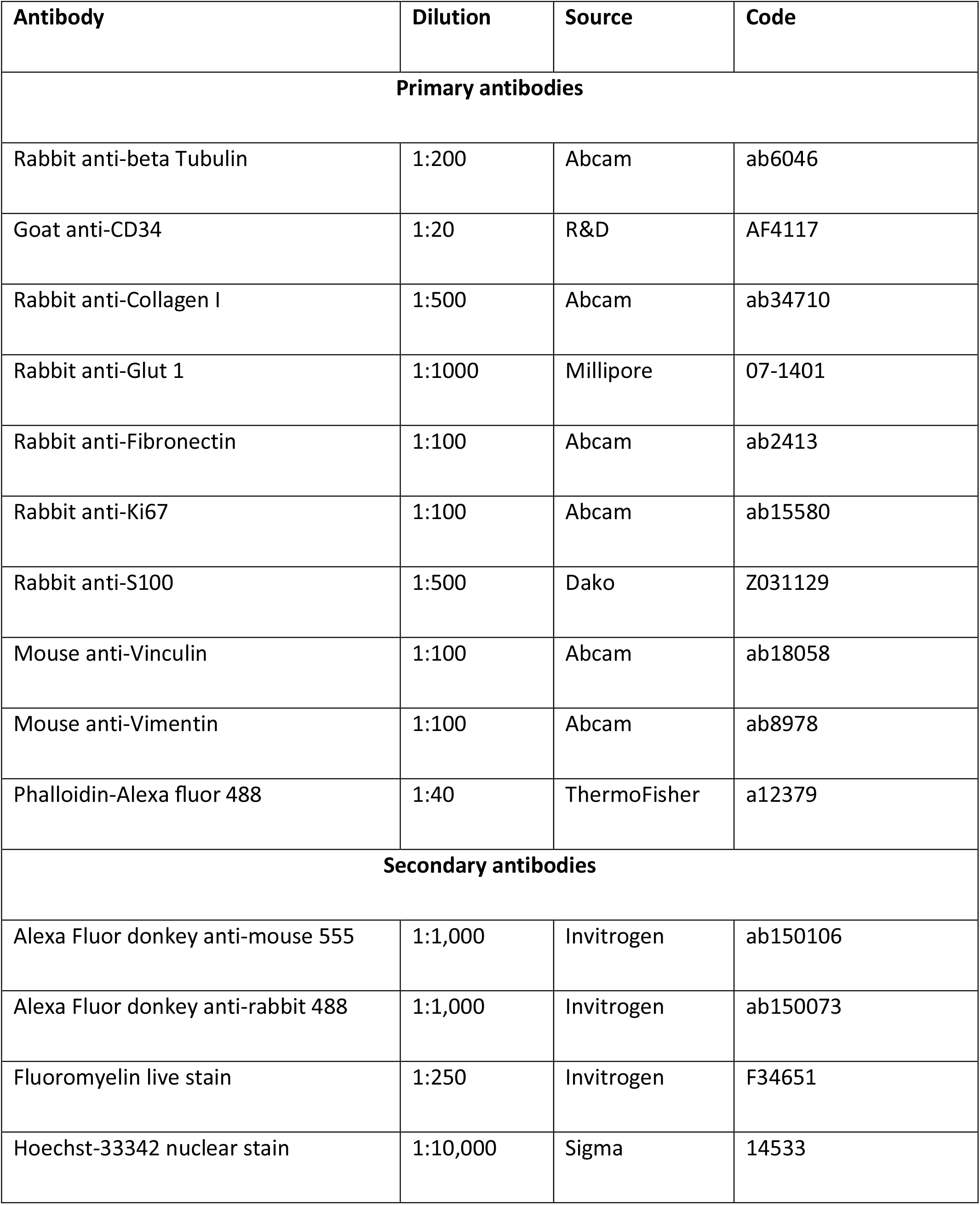
Antibodies used in immunohistochemistry and immunocytochemistry experiments.

## References

1. Chen, Z.-L., Yu, W.-M. & Strickland, S. Peripheral Regeneration. Annu. Rev. Neurosci. 30, 209–233 (2007).

2. Chen, M. S. et al. Nogo-A is a myelin-associated neurite outgrowth inhibitor and an antigen for monoclonal antibody IN-1. Nature 403, 434–439 (2000).

3. Bradbury, E. J. et al. Chondroitinase ABC promotes functional recovery after spinal cord injury. Nature 416, 636–640 (2002).

4. Fawcett, J. W. The Struggle to Make CNS Axons Regenerate: Why Has It Been so Difficult? Neurochem. Res. 45, 144–158 (2020).

5. Moeendarbary, E. et al. The soft mechanical signature of glial scars in the central nervous system. Nat. Commun. 8, ncomms14787 (2017).

6. Saxena, T., Gilbert, J., Stelzner, D. & Hasenwinkel, J. Mechanical characterization of the injured spinal cord after lateral spinal hemisection injury in the rat. J. Neurotrauma 29, 1747–1757 (2012).

7. Möllmert, S. et al. Zebrafish Spinal Cord Repair Is Accompanied by Transient Tissue Stiffening. Biophys. J. 118, 448–463 (2020).

8. Koser, D. E. et al. Mechanosensing is critical for axon growth in the developing brain. Nat. Neurosci. 19, 1592–1598 (2016).

9. Thompson, A. J. et al. Rapid changes in tissue mechanics regulate cell behaviour in the developing embryonic brain. eLife 8, e39356 (2019).

10. Barriga, E. H., Franze, K., Charras, G. & Mayor, R. Tissue stiffening coordinates morphogenesis by triggering collective cell migration in vivo. Nature 554, 523–527 (2018).

11. Bollmann, L. et al. Microglia mechanics: immune activation alters traction forces and durotaxis. Front. Cell. Neurosci. 9, 363 (2015).

12. Moshayedi, P. et al. Mechanosensitivity of astrocytes on optimized polyacrylamide gels analyzed by quantitative morphometry. J. Phys. Condens. Matter Inst. Phys. J. 22, 194114 (2010).

13. Moshayedi, P. et al. The relationship between glial cell mechanosensitivity and foreign body reactions in the central nervous system. Biomaterials 35, 3919–3925 (2014).

14. Carnicer-Lombarte, A. et al. Mechanical matching of implant to host minimises foreign body reaction. bioRxiv 829648 (2019) doi:10.1101/829648.

15. Engler, A. J. et al. Embryonic cardiomyocytes beat best on a matrix with heart-like elasticity: scar-like rigidity inhibits beating. J. Cell Sci. 121, 3794–3802 (2008).

16. Georges, P. C. et al. Increased stiffness of the rat liver precedes matrix deposition: implications for fibrosis. Am. J. Physiol. Gastrointest. Liver Physiol. 293, G1147–1154 (2007).

17. Christ, A. F. et al. Mechanical difference between white and gray matter in the rat cerebellum measured by scanning force microscopy. J. Biomech. 43, 2986–2992 (2010).

18. Koser, D. E., Moeendarbary, E., Hanne, J., Kuerten, S. & Franze, K. CNS Cell Distribution and Axon Orientation Determine Local Spinal Cord Mechanical Properties. Biophys. J. 108, 2137–2147 (2015).

19. Fawcett, J. W. & Keynes, R. J. Peripheral Nerve Regeneration. Annu. Rev. Neurosci. 13, 43–60 (1990).

20. Jessen, K. R. & Mirsky, R. The repair Schwann cell and its function in regenerating nerves. J. Physiol. 594, 3521–3531 (2016).

21. Topp, K. S. & Boyd, B. S. Structure and Biomechanics of Peripheral Nerves: Nerve Responses to Physical Stresses and Implications for Physical Therapist Practice. Phys. Ther. 86, 92–109 (2006).

22. Gautier, H. O. B. et al. Chapter 12 - Atomic force microscopy-based force measurements on animal cells and tissues. in Methods in Cell Biology (ed. Paluch, E. K.) vol. 125 211–235 (Academic Press, 2015).

23. Richard, L., Védrenne, N., Vallat, J.-M. & Funalot, B. Characterization of Endoneurial Fibroblast-like Cells from Human and Rat Peripheral Nerves. J. Histochem. Cytochem. 62, 424–435 (2014).

24. Seddon, H. J. THREE TYPES OF NERVE INJURY. Brain 66, 237–288 (1943).

25. Gordon, T., Sulaiman, O. & Boyd, J. G. Experimental strategies to promote functional recovery after peripheral nerve injuries. J. Peripher. Nerv. Syst. 8, 236–250 (2003).

26. Swift, J. et al. Nuclear Lamin-A Scales with Tissue Stiffness and Enhances Matrix-Directed Differentiation. Science 341, 1240104 (2013).

27. Lefcort, F., Venstrom, K., McDonald, J. A. & Reichardt, L. F. Regulation of expression of fibronectin and its receptor, alpha 5 beta 1, during development and regeneration of peripheral nerve. Development 116, 767–782 (1992).

28. Cattin, A.-L. et al. Macrophage-Induced Blood Vessels Guide Schwann Cell-Mediated Regeneration of Peripheral Nerves. Cell 162, 1127–1139 (2015).

29. Eldridge, C., Bunge, M., Bunge, R. & Wood, P. Differentiation of axon-related Schwann cells in vitro. I. Ascorbic acid regulates basal lamina assembly and myelin formation. J. Cell Biol. 105, 1023–1034 (1987).

30. Arthur-Farraj, P. J. et al. c-Jun Reprograms Schwann Cells of Injured Nerves to Generate a Repair Cell Essential for Regeneration. Neuron 75, 633–647 (2012).

31. Rosso, G., Liashkovich, I., Young, P. & Shahin, V. Nano-scale Biophysical and Structural Investigations on Intact and Neuropathic Nerve Fibers by Simultaneous Combination of Atomic Force and Confocal Microscopy. Front. Mol. Neurosci. 10, (2017).

32. Rosso, G. & Guck, J. Mechanical changes of peripheral nerve tissue microenvironment and their structural basis during development. APL Bioeng. 3, 036107 (2019).

33. Noble, J., Munro, C. A., Prasad, V. S. S. V. & Midha, R. Analysis of Upper and Lower Extremity Peripheral Nerve Injuries in a Population of Patients with Multiple Injuries. J. Trauma Acute Care Surg. 45, 116–122 (1998).

34. Ushiki, T. & Ide, C. 3-Dimensional Organization of the Collagen Fibrils in the Rat Sciatic-Nerve as Revealed by Transmission Electron and Scanning Electron-Microscopy. Cell Tissue Res. 260, 175–184 (1990).

35. Mizisin, A. P. & Weerasuriya, A. Homeostatic regulation of the endoneurial microenvironment during development, aging and in response to trauma, disease and toxic insult. Acta Neuropathol. (Berl.) 121, 291–312 (2011).

36. Reina, M. A., Sala-Blanch, X., Machés, F., Arriazu, R. & Prats-Galino, A. Connective Tissues of Peripheral Nerves. in Hadzic’s Textbook of Regional Anesthesia and Acute Pain Management (ed. Hadzic, A.) (McGraw-Hill Education, 2017).

37. Sunderland, S. The connective tissues of the peripheral nerves. Brain 88, 841–854 (1965).

38. Grant, C. A., Twigg, P. C. & Tobin, D. J. Static and dynamic nanomechanical properties of human skin tissue using atomic force microscopy: effect of scarring in the upper dermis. Acta Biomater. 8, 4123–4129 (2012).

39. Weber, I. P., Yun, S. H., Scarcelli, G. & Franze, K. The role of cell body density in ruminant retina mechanics assessed by atomic force and Brillouin microscopy. Phys. Biol. 14, 065006 (2017).

40. Antonovaite, N., Hulshof, L. A., Hol, E. M., Wadman, W. J. & Iannuzzi, D. Viscoelastic mapping of mouse brain tissue: Relation to structure and age. J. Mech. Behav. Biomed. Mater. 113, 104159 (2021).

41. Xu, Z., Orkwis, J. A., DeVine, B. M. & Harris, G. M. Extracellular matrix cues modulate Schwann cell morphology, proliferation, and protein expression. J. Tissue Eng. Regen. Med. 14, 229–242 (2020).

42. Urbanski, M. M. et al. Myelinating glia differentiation is regulated by extracellular matrix elasticity. Sci. Rep. 6, 33751 (2016).

43. Rosso, G., Liashkovich, I., Young, P., Röhr, D. & Shahin, V. Schwann cells and neurite outgrowth from embryonic dorsal root ganglions are highly mechanosensitive. Nanomedicine Nanotechnol. Biol. Med. 13, 493–501 (2017).

44. Gu, Y. et al. The influence of substrate stiffness on the behavior and functions of Schwann cells in culture. Biomaterials 33, 6672–6681 (2012).

45. Castelnovo, L. F. et al. Schwann cell development, maturation and regeneration: a focus on classic and emerging intracellular signaling pathways. Neural Regen. Res. 12, 1013–1023 (2017).

46. Dupont, S. et al. Role of YAP/TAZ in mechanotransduction. Nature 474, 179–183 (2011).

47. Holtzmann, K. et al. Brain tissue stiffness is a sensitive marker for acidosis. J. Neurosci. Methods 271, 50–54 (2016).

48. Brown, R. et al. Activity-dependent degeneration of axotomized neuromuscular synapses in WldS mice. Neuroscience 290, 300–320 (2015).

49. Hutter, J. & Bechhoefer, J. Calibration of Atomic-Force Microscope Tips. Rev. Sci. Instrum. 64, 1868–1873 (1993).

50. Hertz, H. Ueber die Berührung fester elastischer Körper. J. Für Reine Angew. Math. 92, 156–171 (1882).

51. Lin, D. C., Shreiber, D. I., Dimitriadis, E. K. & Horkay, F. Spherical indentation of soft matter beyond the Hertzian regime: numerical and experimental validation of hyperelastic models. Biomech. Model. Mechanobiol. 8, 345–358 (2009).

52. Boudou, T., Ohayon, J., Picart, C. & Tracqui, P. An extended relationship for the characterization of Young’s modulus and Poisson’s ratio of tunable polyacrylamide gels. Biorheology 43, 721–728 (2006).

53. Brockes, J. P., Fields, K. L. & Raff, M. C. Studies on cultured rat Schwann cells. I. Establishment of purified populations from cultures of peripheral nerve. Brain Res. 165, 105–118 (1979).

54. Tinevez, J.-Y. et al. TrackMate: An open and extensible platform for single-particle tracking. Methods 115, 80–90 (2017).

